# How sequence alignment scores correspond to probability models

**DOI:** 10.1101/580951

**Authors:** Martin C. Frith

## Abstract

Sequence alignment remains fundamental in bioinformatics. Pairwise alignment is traditionally based on ad hoc scores for substitutions, insertions, and deletions, but can also be based on probability models (pair hidden Markov models: PHMMs). PHMMs enable us to: fit the parameters to each kind of data, calculate the reliability of alignment parts, and measure sequence similarity integrated over possible alignments.

This study shows how multiple models correspond to one set of scores. Scores can be converted to probabilities by partition functions with a “temperature” parameter: for any temperature, this corresponds to some PHMM. There is a special class of models with balanced length probability, i.e. no bias towards either longer or shorter alignments. The best way to score alignments and assess their significance depends on the aim: judging whether whole sequences are related versus finding related parts. This clarifies the statistical basis of sequence alignment.

## 1 Introduction

The main way of analyzing nucleotide and protein sequences is by comparing them to related sequences. This is usually done by defining scores for aligned monomers, insertions, and deletions, then finding alignments with maximal total score. An alternative approach is to use a PHMM: a probability model with probabilities for aligned monomers, insertions, and deletions [7]. The probability approach has three major advantages:

- The probabilities can be fitted to each kind of comparison (e.g. error-prone DNA reads versus a genome). This is expected to make the comparisons more accurate.
- We can estimate the reliability of any alignment part, for example, each column (Figure 1). This is useful because alignments often have parts that are uncertain, due to high divergence or repetitive sequence.
- We can measure similarity between two sequences integrated over possible alignments between them. This is expected to detect subtle relationships more powerfully than single optimal alignments [2, 9].

**Figure 1:**
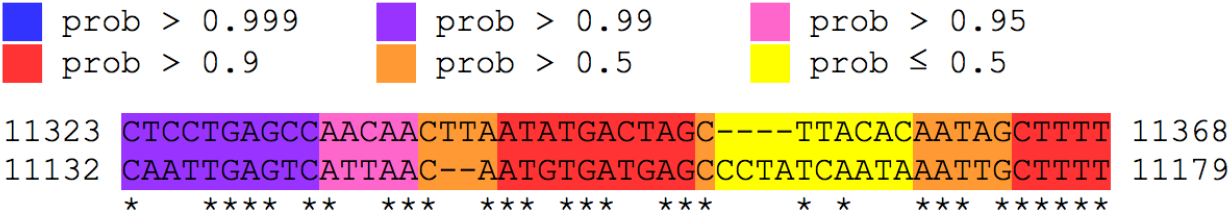
An alignment color-coded by column reliability. The upper sequence is from a human mitochondrial genome, and the lower sequence is from a sea urchin mitochondrial genome. The figure was made using LAST (last.cbrc.jp).

We should bear in mind that “all models are wrong, but some are useful”. Alignment models typically omit rapid evolution of tandem repeats, neighbor-dependence of substitutions, etc., but have proven useful.

Another approach is to define alignment probabilities as exponentiated scores [27, 18]:

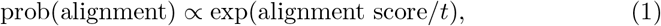

where *t* is a parameter whose value must be chosen somehow. It follows that:

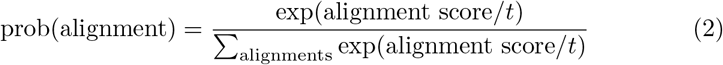

The denominator in Equation (2) is called a “partition function”, and *t* is often called “temperature” due to an analogy with physics. It has been argued that this approach should be distinguished from generative models such as PHMMs [8].

This study describes the equivalence between the partition function approach and PHMMs. The aim is to further unify our understanding of alignment methods, and thereby encourage the use of probability approaches with their aforementioned advantages. This study also clarifies the notion of alignment models with balanced length probability, i.e. no bias towards either longer or shorter alignments. It concludes with a discussion of the best way to score alignments and assess their significance, depending on our precise aim. One previous study also describes a one-to-many relationship between alignment scores and probabilities [1], but lacks most of the results presented here.

### 1.1 Review of score-based alignment

We wish to align two sequences:

*R*_1_,…, *R_m_*: 1st sequence (e.g. “reference”), of length *m*.
*Q*_1_,…, *Q_n_*: 2nd sequence (e.g. “query”), of length *n*.

The classic approach is to specify a scoring scheme, which assigns a numeric score for aligning any pair of letters or inserting a gap:

*S_xy_*: score for aligning reference letter *x* to query letter *y*.
*a_D_*: first deletion score.
*b_D_*: deletion extension score. A deletion of length *k* scores *a_D_* + *b_D_* × (*k* – 1).
*a_I_*: first insertion score.
*b_I_*: insertion extension score. An insertion of length k scores *a_I_* + *b_I_* × (*k* – 1).

This describes *affine* gap scores, proportional to the gap length plus a constant.^1^ This is not the only way to score gaps, but it is the most common. Usually, *a_D_* ≤ *b_D_* ≤ 0 and *a_I_* ≤ *b_I_* < 0. Having defined a scoring scheme, we seek alignments with maximal total score.^2^ Sometimes we seek an optimal *global* alignment (which aligns the sequences end-to-end), sometimes a *local* alignment (which aligns a pair of substrings).

#### 1.1.1 Degrees of freedom

Some transformations of the score parameters have no effect on alignment. For local alignment, these transformations have one degree of freedom: if we multiply all the scores by any positive constant there is no effect on alignment. For global alignment, there are two further degrees of freedom. First, we can add any constant (say h) to *a_D_, b_D_* and *S_xy_*: this also has no effect on alignment, because it simply adds *h* × *m* to the total score of any alignment. Second, we can add any constant to *a_I_*, *b_I_* and *S_xy_*. These freedoms can be used to set *b_D_* = *b_I_* = 0, which might allow global alignment software to run a bit faster [17].

#### 1.1.2 Algorithms for local alignment

Maximal-scoring local alignments can be found by a classic algorithm, which builds up the solution using prefixes of the sequences [22, 16]. This algorithm has several variants.

One variant calculates the optimal score for aligning the length-*i* prefix of *R* to the length-*j* prefix of *Q*, ending with *R_i_* aligned to *Q_j_* (*X_ij_*), *R_i_* aligned to a gap (*Y_ij_*), or *Q_j_* aligned to a gap (*Z_ij_*):

**Algorithm I**

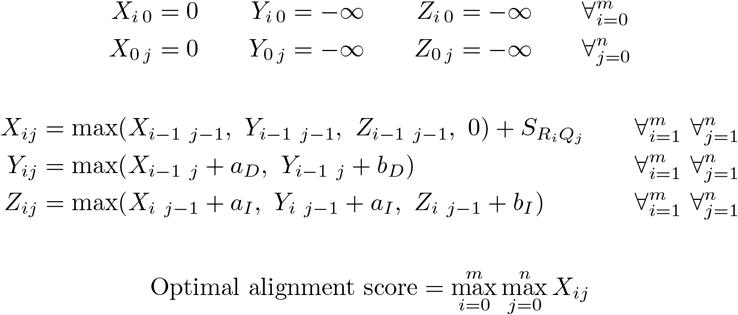

This variant is asymmetric in that it considers insertions after deletions but not deletions after insertions. This is because those two configurations always have the same score, so we may as well only consider the former. The algorithm only returns the alignment score, but an actual alignment can be found by tracing back from a maximal-scoring (*i,j*) endpoint.

Another variant uses *W_ij_* = max(*X_ij_*, *Y_ij_*, *Z_ij_*, 0) instead of *X_ij_*:

**Algorithm II**

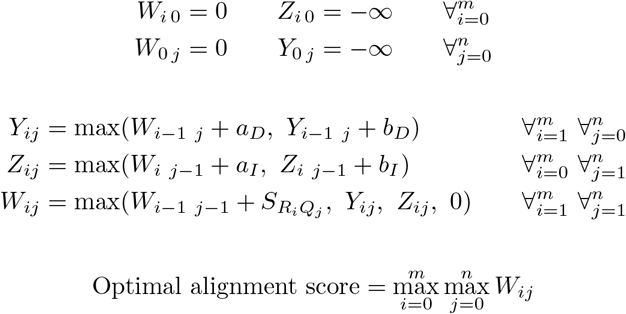

This variant gives identical results to Algorithm I (assuming that *a_D_ ≤ b_D_* and *a_I_* ≤ *b_I_*), but is more efficient (fewer operations). There are even more efficient variants [29, 5, 10, 21, 23], discussed in the Supplement.

### 1.2 Review of alignment probability models

A pair hidden Markov model (PHMM) is a scheme for randomly generating a pair of sequences. Let us first consider a gapless model, which is simpler but illustrates the main issues (Figure 2). Starting at “begin”, we randomly traverse the arrows until we hit the “end”. Whenever we have a choice of arrows, we randomly choose one, with the labeled probabilities: for example, at the first choice point we choose one arrow with probability *ω_D_* and the other with probability 1 – *ω_D_*. Whenever we reach a circle labeled “D”, we randomly generate a reference sequence letter, with probabilities *ϕ* (e.g. *ϕ_a_* = 0.3, *ϕ_c_* = 0.2, *ϕ_g_* = 0.2, *ϕ_t_* = 0.3). Likewise, at “I” we generate a query letter, and at “M” we generate an aligned pair of letters. Thus, the model generates zero or more reference letters, then zero or more query letters, then zero or more aligned letter pairs, then zero or more reference letters, and finally zero or more query letters: a gapless local alignment.

**Figure 2:**
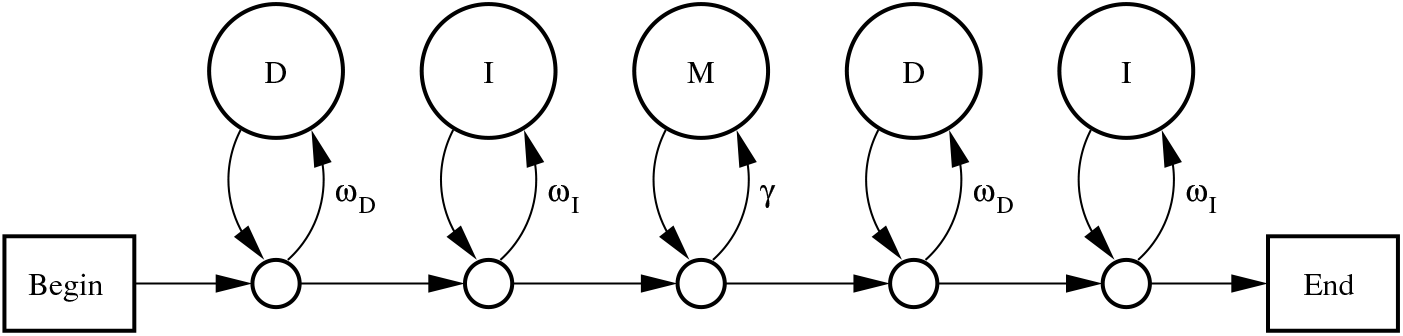
A PHMM for gapless local alignment. The state labeled M emits
aligned letters *x*: *y* with probability *π_xy_*. States labeled D emit reference letters *x* with probability *ϕ_x_*. States labeled I emit query letters *y* with probability *ψ_y_*.

Typically, a PHMM is used not to generate but to analyze a pre-existing pair of sequences. The simplest analysis is to find a path through the model (i.e. an alignment) most likely to have generated those sequences. Consider an alignment where the first *c* letters of *R* and *d* letters of *Q* are unaligned, the next *e* letters of *R* and *Q* are aligned, and the final *f* letters of *R* and *g* letters of *Q* are unaligned. The probability of this alignment under the model is:

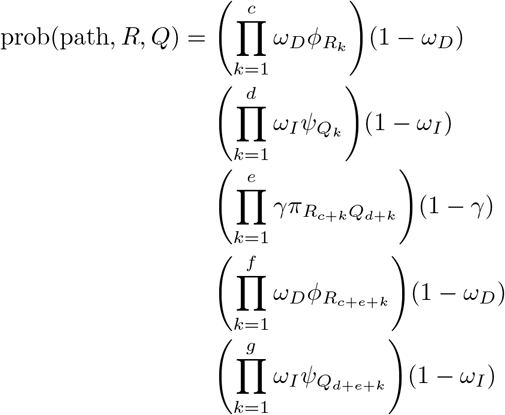

This can be simplified by factoring out a constant *μ*, defined as:

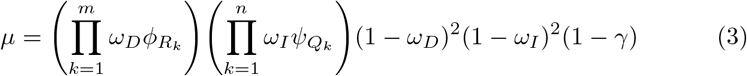

Because *μ* does not depend on the path, we can find a most-probable path by maximizing:

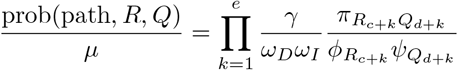

Finally, because maximizing a value is equivalent to maximizing its logarithm, we can maximize:

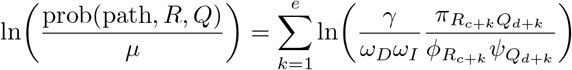

This reveals the connection between model-based and score-based alignment. If we define a substitution score matrix as follows:

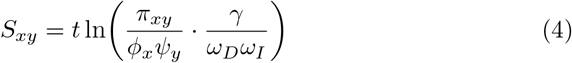

then a most-probable path is one which maximizes the sum of scores. Note that *μ* is the probability of a null alignment (with zero aligned letters): so an alignment score is a log probability ratio relative to a null alignment.

## 2 Degrees of freedom in the gapless model

The model in Figure 2 has several degrees of freedom, meaning that we can vary the parameters with no effect on *S_xy_* and thus no effect on alignment.

First, we can freely vary *ω_D_, ω_I_*, and *γ*, provided that 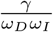 remains fixed. This is rather interesting: it means, for example, that a PHMM with *ω_D_* = 0.8, *ω_I_* = 0.2 is equivalent to one with *ω_D_* = *ω_I_* = 0.4. These PHMMs would not be equivalent if used to generate aligned sequences: the former would tend to produce lopsided pairs with query shorter than reference sequence. When used to analyze pre-existing sequences, though, the sequence lengths are fixed in advance, and the two models become equivalent.

Second, we can freely vary *ϕ, ψ*, and *t*, provided that we co-vary *π* and 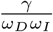 in a suitable way. If we fix values for *S, ϕ, ψ*, and *t*; then *π* and 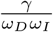 are given by this equation, where the right-hand side is fully specified:

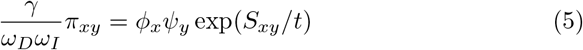

Since *π_xy_* must sum to 1, we can infer that:

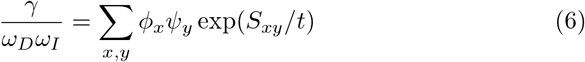

Then we can calculate *π* using Equation (5).

### 2.1 Homogeneous letter probabilities

The letter probabilities in aligned and unaligned regions need not be the same, but we may wish to make them so [26]:

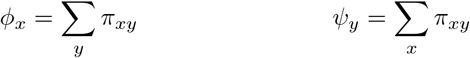

In this case, we can freely vary *t*, provided we co-vary *π* and 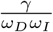 in a suitable way. If we fix values for *S* and *t*, and define 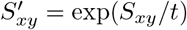 and 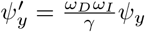, we get:

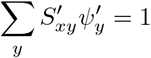

This is a set of simultaneous linear equations, which can be solved by standard methods (provided that *S*′ is not a singular matrix). We can then calculate:

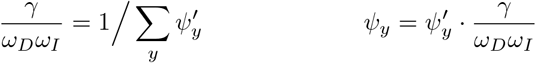

*ϕ* can be calculated in the same way. These calculations may yield negative *ϕ_x_* or *ψ_y_*, meaning that these values of *S* and *t* are not consistent with homogeneous letter probabilities.

### 2.2 Uniform length probability

We may wish to focus on the case where 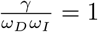, which means that alignments of different lengths have uniform prior probability. (Hidden Markov models are sometimes criticized for having geometrically decaying length distributions [8]: this is true when a model is used to generate sequences, but not necessarily when it is used to analyze pre-existing sequences, as in this case where we have uniform length probability.)

If we require both uniform length probability and homogeneous letter probabilities, *π* has no degrees of freedom (usually). Starting from a given substitution score matrix *S, t* must have a value such that 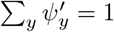. Yu et al. showed that *t* has at most one such value that also yields *ϕ* and *ψ* ≥ 0, and they described a numerical procedure for finding it [26, 24]. It is possible (though unlikely) to have a solution where *S*’ is singular, in which case *π* does have degrees of freedom, i.e. a linear space of values that yield the same *S* [24].

## 3 Examples

For brevity, let us consider DNA models, although the theory applies equally to proteins. A substitution score matrix named “HoxD70” is often used for inter-species genome alignment [6]:

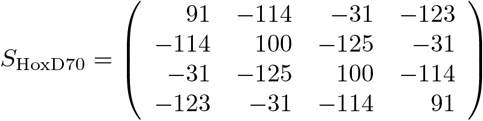

Assuming a PHMM of the form shown in Figure 2, with homogeneous letter probabilities, any of the parameter combinations in Table 1 yield HoxD70. This is rather striking: apparently-different PHMM parameter settings lead to the exact same score parameters, and therefore always produce the same optimal alignments.

**Table 1:** Sets of PHMM parameters that correspond to the HoxD70 score matrix.

| $t$ | $\frac{\gamma}{\omega_D \omega_I}$ | $\phi = \psi$ | $\pi$ |
| --- | --- | --- | --- |
| 30 | 6.06 | $\begin{pmatrix} 0.288 \\ 0.212 \\ 0.212 \\ 0.288 \end{pmatrix}$ | $\begin{pmatrix} 0.284 & 0.000225 & 0.00359 & 0.000226 \\ 0.000225 & 0.208 & 0.000115 & 0.00359 \\ 0.00359 & 0.000115 & 0.208 & 0.000225 \\ 0.000226 & 0.00359 & 0.000225 & 0.284 \end{pmatrix}$ |
| 96.1735 | 1 | $\begin{pmatrix} 0.266 \\ 0.234 \\ 0.234 \\ 0.266 \end{pmatrix}$ | $\begin{pmatrix} 0.182 & 0.019 & 0.0451 & 0.0197 \\ 0.019 & 0.155 & 0.0149 & 0.0451 \\ 0.0451 & 0.0149 & 0.155 & 0.019 \\ 0.0197 & 0.0451 & 0.019 & 0.182 \end{pmatrix}$ |
| 10000 | 0.996 | $\begin{pmatrix} 0.258 \\ 0.242 \\ 0.242 \\ 0.258 \end{pmatrix}$ | $\begin{pmatrix} 0.0673 & 0.062 & 0.0625 & 0.0658 \\ 0.062 & 0.0596 & 0.0583 & 0.0625 \\ 0.0625 & 0.0583 & 0.0596 & 0.062 \\ 0.0658 & 0.0625 & 0.062 & 0.0673 \end{pmatrix}$ |

The “Simple” substitution score matrix provides another example:

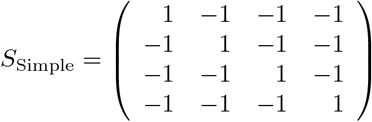

Any of the PHMM parameter sets in Table 2 yield this matrix (again using homogeneous letter probabilities).

**Table 2:** Sets of PHMM parameters that correspond to the Simple score matrix.

| $t$ | $\frac{\gamma}{\omega_D \omega_I}$ | $\phi = \psi$ | $\pi$ |
| --- | --- | --- | --- |
| 0.3 | 7.03 | $\begin{pmatrix} 0.25 \\ 0.25 \\ 0.25 \\ 0.25 \end{pmatrix}$ | $\begin{pmatrix} 0.249 & 0.000317 & 0.000317 & 0.000317 \\ 0.000317 & 0.249 & 0.000317 & 0.000317 \\ 0.000317 & 0.000317 & 0.249 & 0.000317 \\ 0.000317 & 0.000317 & 0.000317 & 0.249 \end{pmatrix}$ |
| 0.910239 | 1 | $\begin{pmatrix} 0.25 \\ 0.25 \\ 0.25 \\ 0.25 \end{pmatrix}$ | $\begin{pmatrix} 0.188 & 0.0208 & 0.0208 & 0.0208 \\ 0.0208 & 0.188 & 0.0208 & 0.0208 \\ 0.0208 & 0.0208 & 0.188 & 0.0208 \\ 0.0208 & 0.0208 & 0.0208 & 0.188 \end{pmatrix}$ |
| 10 | 0.955 | $\begin{pmatrix} 0.25 \\ 0.25 \\ 0.25 \\ 0.25 \end{pmatrix}$ | $\begin{pmatrix} 0.0723 & 0.0592 & 0.0592 & 0.0592 \\ 0.0592 & 0.0723 & 0.0592 & 0.0592 \\ 0.0592 & 0.0592 & 0.0723 & 0.0592 \\ 0.0592 & 0.0592 & 0.0592 & 0.0723 \end{pmatrix}$ |

These examples show a common pattern. When *t* is low, 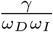 is high, and *π* has low mismatch probabilities. High 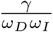 means that longer alignments are preferred, which counteracts the aversion to mismatches. As *t* increases, 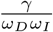 decreases, and the mismatch probabilities increase: the net effect is that optimal alignments remain the same.

As *t* increases further, however, 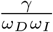 reaches a minimum (whose value differs slightly in the two examples), and then rises asymptotically to 1 (Figure 3). This means that not any value of 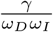 can be consistent with a given substitution score matrix: the value must be ≥ this minimum.

**Figure 3:**
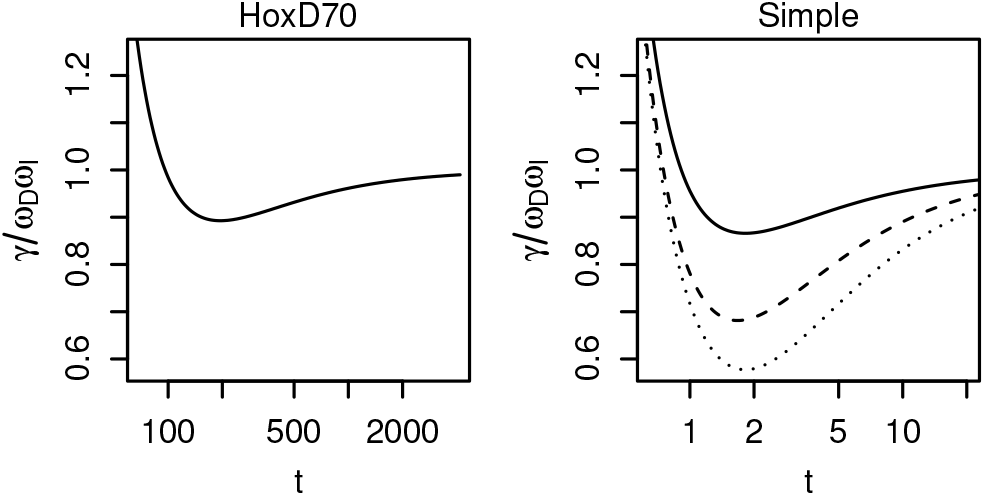
How 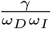 varies with *t* for the HoxD70 matrix and the Simple matrix. Dashed line: effect of changing the mismatch score to −2. Dotted line: effect of changing the mismatch score to –3.

### 3.1 Effect of mismatch score

If we strengthen the Simple matrix’s mismatch score (i.e. make it more negative), the value of 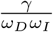 decreases for any given *t* (Figure 3, Equation 6). One consequence is that uniform length probability occurs at lower values of *t*.

## 4 Linear gap costs

Let us now consider a simple gapped model (Figure 4). Just as for the gapless model, we can align two sequences by finding a path through the model most likely to have generated the sequences. This is equivalent to maximizing an alignment score defined as *t* ln(prob(path, *R, Q*)/*μ_G_*), where *μ_G_* is the probability of a null alignment (that never traverses the *γ*, *α_D_* or *α_I_* arrows):

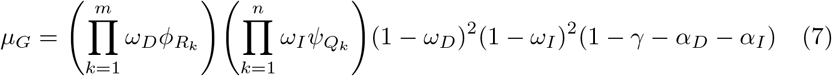

**Figure 4.**
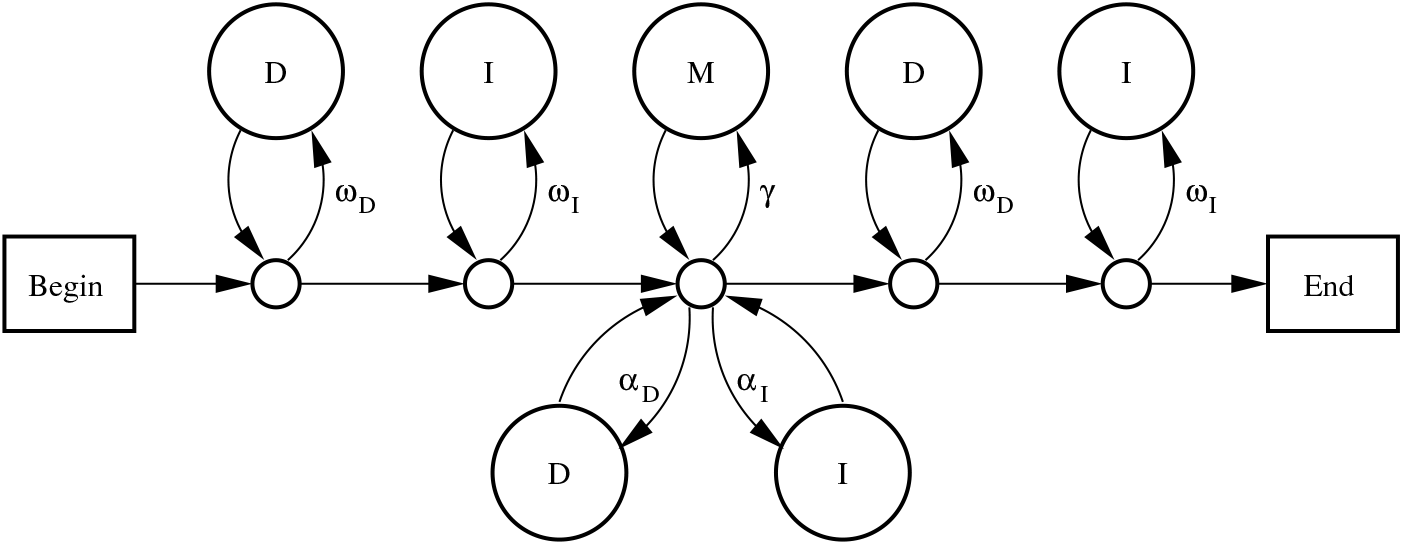
A simple PHMM for gapped local alignment. The state labeled M emits aligned letters *x: y* with probability *π_xy_*. States labeled D emit reference letters *x* with probability *ϕ_x_*. States labeled I emit query letters *y* with probability *ψ_y_*.

This alignment score is a sum of scores for substitutions, deletions, and insertions:

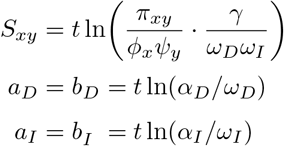

### 4.1 Degrees of freedom

This model has several degrees of freedom, meaning that we can vary the model probabilities with no effect on alignment scores. First, we can freely vary *ω_D_* and *ω_I_*, provided we co-vary *γ, α_D_*, and *α_I_* so as to keep these values fixed:

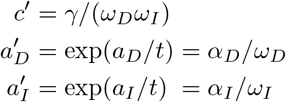

Second, we can freely vary *ϕ, ψ*, and *t*, provided we co-vary *π, c′, α_D_/ω_D_*, and *α_I_/ω_I_* in a suitable way. Overall, the degrees of freedom are similar to those of the gapless model.

### 4.2 Balanced length probability

There is a useful gapped generalization of gapless alignment with uniform length probability. If *ω_D_* and *ω_I_* are relatively low, but *γ, α_D_*, and *α_I_* are high, the model has a bias towards longer alignments. Conversely, if *ω_D_* and *ω_I_* are high but *γ, α_D_*, and *α_I_* are low, the model is biased towards shorter alignments. A natural balance occurs when:

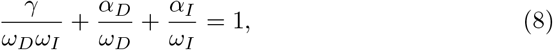

or equivalently:

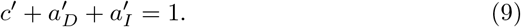

If 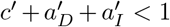, the degrees of freedom allow us to vary *ω_D_* and *ω_I_* arbitrarily close to 1, in which case *γ* + *α_D_* + *α_I_* (the probability of continuing the alignment) will be less than 1. This means there is a bias towards shorter alignments. Conversely, if 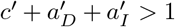, we cannot vary both *ω_D_* and *ω_I_* arbitrarily close to 1: they have an upper bound that occurs when *γ* + *α_D_* + *α_I_* = 1. This means there is a bias towards longer alignments.

Balanced length probability can also be understood as conservation of probability ratio relative to a null alignment (see the Supplement and [25]).

## 5 Affine gap costs

There is more than one PHMM topology corresponding to affine gap alignment: Figure 5 shows two options. Model A is more “ambiguous”, meaning that different paths through the PHMM yield indistinguishable alignments. For example, if we have an alignment with two consecutive deleted bases, this could be one deletion of length two (traversing the arrow labeled *β_D_*), or two deletions of length one (traversing *α_D_* twice). Using model B, on the other hand, it would have to be one deletion of length two. Model A is arguably more realistic, because in real evolution two independent, consecutive deletions might occur. I believe model A is the most elegant possible model for local affine-gap alignment: it makes insertions and deletions symmetric, and produces simple equations.

**Figure 5:**
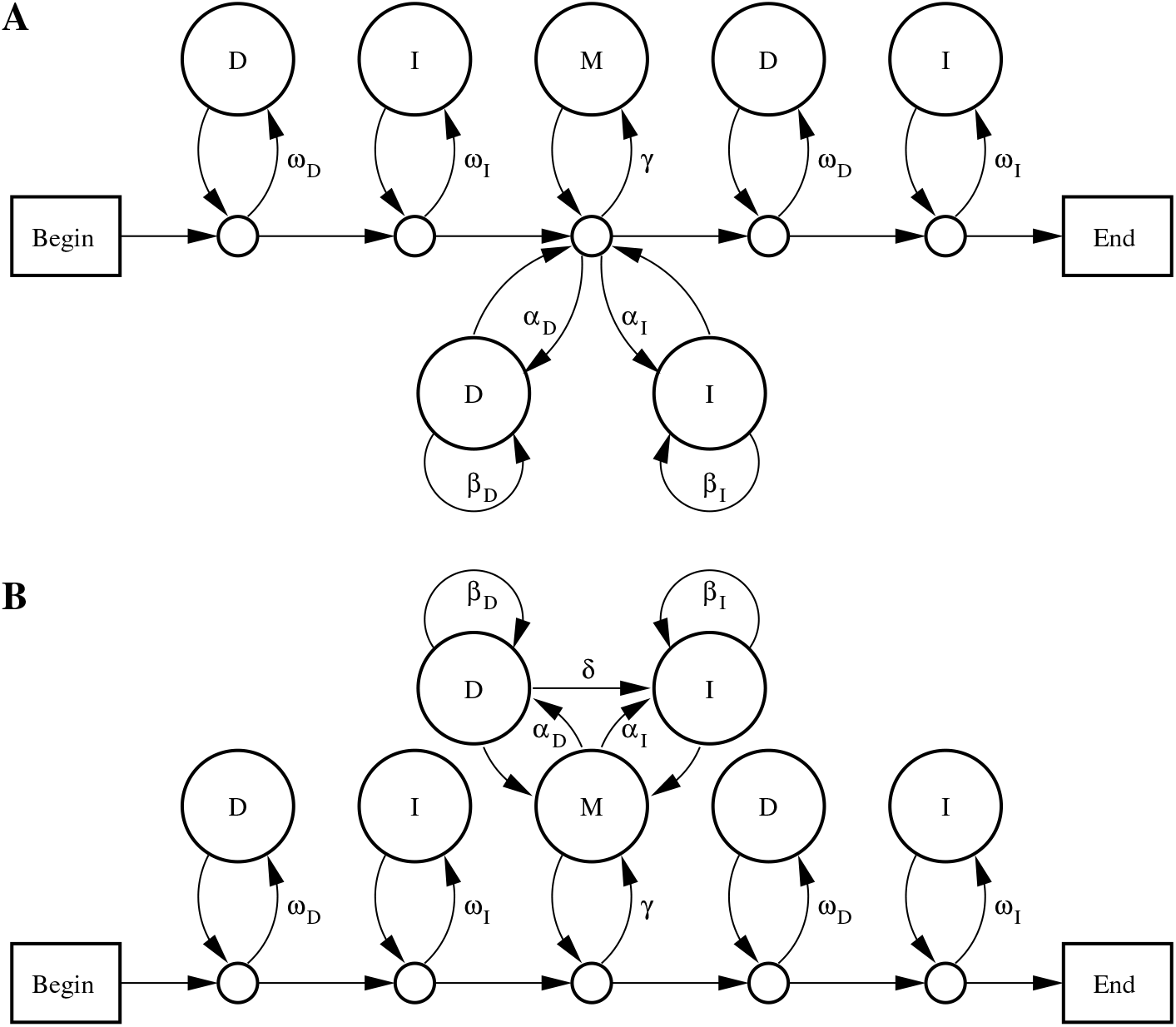
Two PHMMs for gapped local alignment. States labeled M emit aligned letters *x: y* with probability *π_xy_* States labeled D emit reference letters *x* with probability *ϕ_x_* States labeled I emit query letters *y* with probability *ψ_y_*.

Let us now relate model A probabilities to alignment scores. We must first decide whether “alignment” means “path” or “set of indistinguishable paths”: let us use the former definition here, and explore the latter in the Supplement. An alignment score is then *t* ln(prob(path, *R, Q*)/*μ_G_*), which is a sum of scores for substitutions, deletions, and insertions:

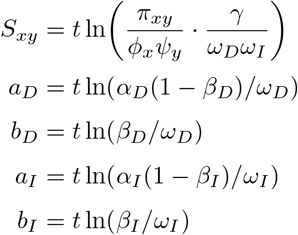

### 5.1 Degrees of freedom

The affine-gap models have several degrees of freedom, meaning that we can vary the model probabilities with no effect on alignment scores. In model A we can freely vary *ω_D_* and *ω_I_*, provided we co-vary *β_D_, β_I_*, *α_D_*, *α_I_*, and *γ* so as to keep these values fixed:

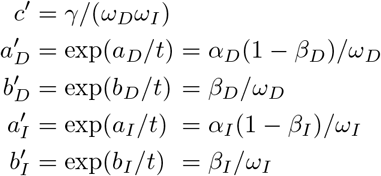

There are also degrees of freedom involving *ϕ, ψ*, and *t*, similar to the gapless and linear-gap models.

### 5.2 Limits to degrees of freedom

Our freedom to vary *ω_D_* and *ω_I_* (while keeping *c*′ etc. fixed) may have an upper limit. In model A, we must have *γ + α_D_ + α_I_* < 1, and:

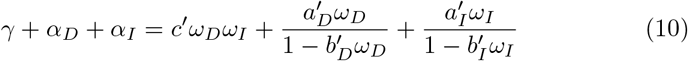

Thus, *γ + α_D_ + α_I_* increases with increasing *ω_D_* and *ω_I_*. (Let us assume that *b_D_* and *b_I_* are negative, thus 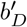 and 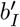 are < 1, thus the denominators in Equation (10) are always positive.) This may prevent high values of *ω_D_* and *ω_I_*. Conversely, *γ + α_D_ + α_I_* may have an upper limit: its maximum possible value occurs when *ω_D_* ≈ 1 ≈ *ω_I_*.

Let us see some examples (Figure 6), with symmetric insertions and deletions, so we can drop the *D* and *I* subscripts (e.g. *ω_D_* = *ω_I_* = *ω*). The upper limit on *ω* can be found by solving this cubic equation:

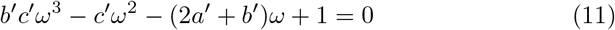

**Figure 6:**
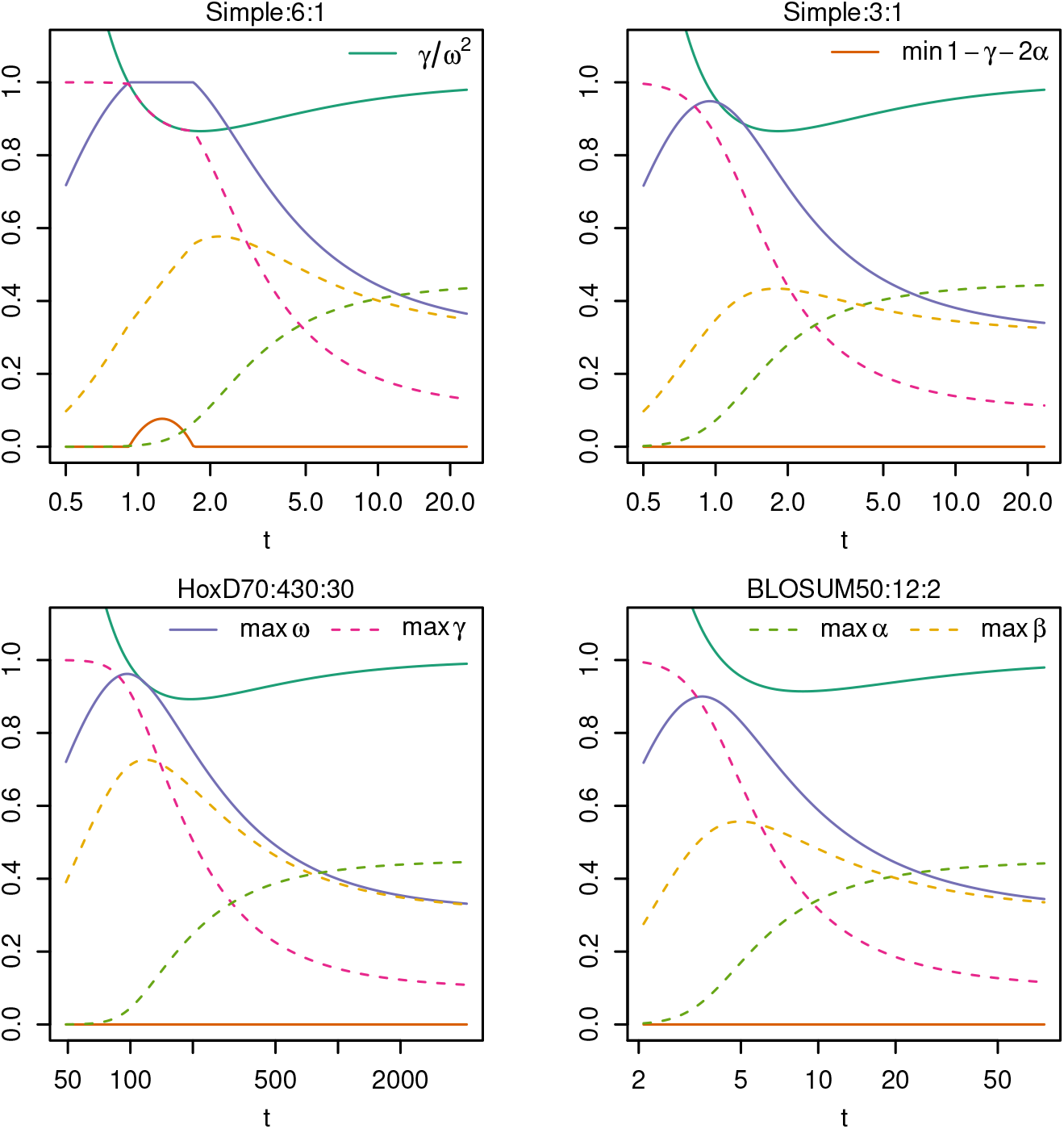
How model A’s parameter limits vary with *t*, for four scoring schemes. Each scoring scheme is written as three colon-separated components, matrix: first gap cost: gap extension cost.

These examples assume homogeneous letter probabilities, so that *c*′ = *γ/ω*^2^ is a unique function of *t*. In all cases, as *t* increases, the maximum possible value of *ω* rises to a peak, and then decreases towards 0.311. For the Simple:6:1 scoring scheme, there is a range of *t* for which *ω* can be arbitrarily close to 1. In precisely this range, *γ* + 2*α* (i.e. *γ + α_D_ + α_I_*) cannot be arbitrarily close to 1. Both can be arbitrarily close to 1 at the two endpoints of this range.

For the other scoring schemes, *ω* can never be arbitrarily close to 1. Its maximum possible value is 0.948 for Simple:3:1 (when *c*′ ≈ 0.980), 0.962 for HoxD70:430:30 (when *c*′ ≈ 0.999), and 0.900 for BL0SUM50:12:2 (when *c*′ ≈ 1.08).

The other parameters have limits too. When *t* is low *γ* can be arbitrarily close to 1, but as *t* increases the maximum possible *γ* steadily decreases towards 0.311^2^. As *t* increases, the maximum possible *β* rises to a peak, before decreasing towards 0.311. The maximum possible *α* is extremely low when *t* is low, but rises towards 0.452.

Previously, Durbin et al. reverse-engineered a PHMM for BLOSUM50:12:2 [7, Section 4.5]. Their PHMM has *ω* = 0.8 and *γ* = 0.64 (though its topology is slightly different from model A), which they argue is unrealistic since it implies very short sequences. The reverse-engineering problem is however underdetermined: e.g. when *c*′ = 1, the maximum possible *ω* is 0.88 and the maximum possible γ is 0.77. This probably does not alter their conclusion.

### 5.3 Balanced length probability

For model A, balanced length probability occurs when:

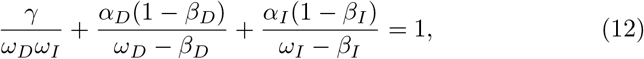

or equivalently:

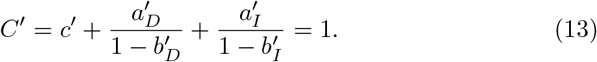

When *C*′ < 1, there is an upper limit on *γ* + *α_D_* + *α_I_* (Equation 10). When *C*′ > 1, there is an upper limit on *ω_D_* and *ω_I_*. Only when *C*′ =1 are there no such limits.

### 5.4 Non-uniqueness of *t*

For gapped (unlike gapless) alignment, balanced length probability does not imply a unique value for *t*. Recall that, if we assume homogeneous letter probabilities, there is one degree of freedom involving *π, ϕ, ψ, c*′, and *t*. If we wish *C*′ =1, then *c*′ must have some value < 1 (Equation 13, assuming 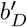 and 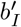 are < 1). Figure 3 shows there are two values of *t* for *c*′ slightly less than 1, and no *t* for *c*′ much less than 1. Accordingly, Figure 6 shows balanced length probability at two values of *t* for the Simple:6:1 scoring scheme, and none for the other three schemes. If the gap and mismatch scores are strong (highly negative), balanced length probability implies *c*′ slightly less than 1, thus two ts. If they are weak (near zero), balanced length probability implies *c*′ much less than 1, which never occurs for any *t*.

## 6 Sum of alignment probabilities

Probability models enable various useful calculations (e.g. Figure 1), which are based on the total probability of all possible alignments [7]:

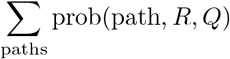

This can be calculated by the Forward algorithm [7]. The Forward algorithm for model A uses these recurrence relations, which calculate the sum of probabilities for aligning the length-*i* prefix of *R* with the length-*j* prefix of *Q* 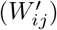, ending with *R_i_* aligned to *Q_j_* 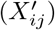, *R_i_* aligned to a gap 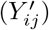, or *Q_j_* aligned to a gap 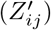:

**Algorithm III**

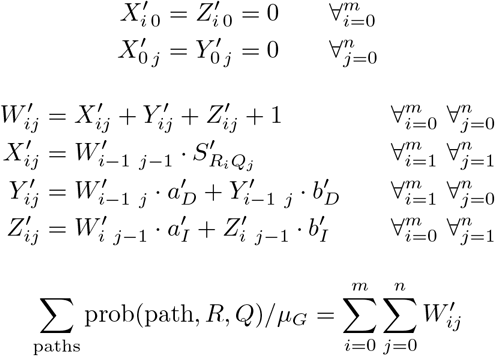

This algorithm uses four dynamic programming matrices, but *X*′ could easily be eliminated (at the expense of making the initializations a bit less simple).

Algorithm III is almost a mechanical transformation of Algorithm II. Firstly, since it uses probabilities without log transformation, addition is replaced by multiplication, and the primitive terms are replaced by exp(•/*t*). Secondly, since it calculates the sum of probabilities rather than the optimal probability, maximization is replaced by summation. The only nuance is that the initialization of *W*′ has to be specified more carefully.

The Forward algorithm for model B, on the other hand, is very similar to Algorithm I (see the Supplement). The important point is that, although Algorithms I and II give identical results, the two Forward algorithms do not. The precise form of the algorithm reflects the model topology, i.e. which possible paths there are. To find optimal alignments, this need not be fully specified, because there are paths that cannot be optimal (e.g. two length-1 deletions in a row), and so we need not care whether the model allows them. To calculate the total probability, on the other hand, we do need to specify the topology (possible paths).

## 7 Discussion

This study describes the many-to-one relationship between probability models and score parameters for sequence alignment. In order to perform alignment probability calculations with the Forward algorithm, we need not fully specify the probability model. Starting from an alignment scoring scheme, we need only specify two further things: (i) a value for *t*, and (ii) a model topology (or equivalently, the form of the dynamic programming algorithm). Typically, any value of *t* is valid and corresponds to some PHMM.

In practice, alignment is often done with ad hoc gap costs, but a probability-based score matrix (like HoxD70 or BLOSUM62) with homogeneous letter probabilities and uniform (gapless) length probability. For such a score matrix, it seems reasonable to use the unique *t* that recovers the original letter probabilities (*π_xy_*). For gapped alignment, however, this corresponds to a model without balanced length probability: weaker gap costs increasingly favor longer alignments. This suggests that such scoring schemes are “wrong”, and should be fixed by subtracting a value (e.g. *t* ln[*ω_D_ω_I_*/*γ*]) from the matrix scores.

On the other hand, if we get an alignment scoring scheme from a model with homogeneous letter probabilities and balanced length probability, it is generally impossible to recover the original *t*, because it is not unique.

### 7.1 Useful probability calculations

This study highlights the importance of Formula (1), which is fairly simple and should be more widely known. Although this formula is true by fiat in the partition function approach, it is true by derivation in the PHMM approach, provided we use the *t* corresponding to our PHMM. Even if we use the “wrong” *t*, it will correspond to some other PHMM.

An example of the formula’s utility arises in alignment of short DNA reads to a reference genome. Suppose one read aligns strongly to three loci, *A, B*, and *C*, with scores *s_A_, s_B_*, and *s_C_*. The probability that A is correct is:

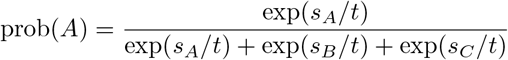

This assumes that exactly one locus is correct, which would be a good assumption for an *ancestral* reference, without reference-specific deletions or duplications [13]. Ideally the denominator would sum over all possible alignments, and perhaps the numerator would sum over alternative alignments to the same locus, but in practice this simple calculation often works well [15]. It is worth emphasizing the generality of this calculation, e.g. it remains valid if we use specialized alignment parameters to model AT-rich genomes or bisulfite-converted DNA [14], or incorporate sequence quality data into the model [15].

A more sophisticated use of the formula is to estimate the probability that an alignment part is correct:

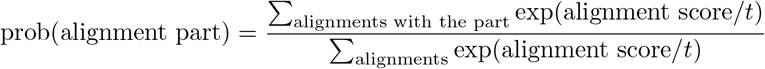

In particular, we can estimate the probability that each column of an alignment is correct (Figure 1) [7]. This is important when studying sequence divergence at fine resolution.

### 7.2 Sequences with multiple similar segments

The models in this study (Figure 2, 4, 5) model sequences with *one* similar segment, but long sequences often have multiple similar segments. This particular mismatch between model and reality can cause standard sequence comparison methods to work poorly [28]. This can be addressed by modeling a *set* of alignments [12] (which is more tractable when comparing a derived sequence to an ancestral sequence [13]): but this method uses a partition function approach without clear correspondence to a generative probability model.

### 7.3 Alignment significance

A fundamental task is detection of significant sequence similarities, i.e. similarities stronger than is likely to occur by chance. “By chance” is usually defined as: between random sequences of independent monomers. “Strength” is usually defined as optimal alignment score, though a more powerful definition can be made by integrating over alternative alignments [2, 9].

Optimal local alignment scores of random sequences typically follow a Gumbel distribution, enabling us to know the significance of any score. For gapped alignment, it is not known how to determine this distribution from first principles, but typically it can be determined rapidly by importance sampling [20]. There is evidence that significant similarity reliably indicates homology of biological sequences, only if simple repeats are excluded in a particular way [11].

There is a special kind of alignment for which scores of random sequences appear to follow a simple distribution [25]. Here, alignment scores are computed by a modified Forward algorithm, where the final summation is replaced by maximization (e.g. 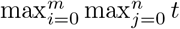 ln 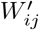 in Algorithm III). Thus, these scores integrate over some, but not all, alternative alignments. (Note that integrating over all alignments is not necessarily desirable. E.g. if we compare the human and mouse X chromosomes, our aim is probably not to judge whether the whole chromosomes are related.) If we use parameters with balanced length probability, these alignment scores follow a Gumbel distribution with scale parameter *λ* = 1/*t*. This is another reason to favor models with balanced length probability.

### 7.4 Aims of sequence comparison

One possible aim of sequence comparison is to judge whether two given sequences are related. For this aim, it is appropriate to calculate a likelihood ratio, for models of related and unrelated sequences [19, 8]. Typical alignment methods do not do this, e.g. the Forward algorithm in this study outputs a probability ratio whose denominator is not the likelihood of an unrelated-sequences model. One symptom of this is that optimal alignment scores between random sequences tend to increase with sequence length [8].

A different aim, more relevant for longer sequences, is to find related segments. For example, if we compare the human and mouse X chromosomes, it is of limited use to obtain one likelihood ratio indicating whether these chromosomes are related. We probably wish to find related parts. For this aim, it is natural that optimal alignment scores between random sequences tend to increase with sequence length, because the search space increases.

A related issue is how to report the significance of sequence similarities. A BLAST *E*-value is: the expected number of “distinct” alignments with greater or equal score, between two random sequences with lengths equal to the given query sequence and the database [4, 3]. This means that *identical* alignments have *different E*-values depending on the query length. For example, if we find three identical alignments, two in a long query (say chromosome 1) and one in a short query (say chromosome Y), only the latter may be deemed significant. This is appropriate for judging whether each whole query has a significant match, but not for finding significant matches in a set of queries. For the latter aim, significance could be reported as, say, expected number of alignments per million query bases.

## 8 Acknowledgements

I thank John Spouge, Michiaki Hamada, Kiyoshi Asai, Anish Shrestha, and members of the CBRC Genome Meeting for useful comments.

## Footnotes

1 There are two common parameterizations of affine score for a gap of length *k*: *a* + *b* × *k* and *a* + *b* × (*k* – 1). It turns out the latter is a better fit to the mathematics of this study.

2 In the unusual case that *a_D_* > *b_D_* or *a_I_* > *b_I_*, we must decide whether a gap of length *k* may be scored as *k* separate gaps. Algorithm I does not allow this; Algorithm II does.

