## Supplement for "How sequence alignment scores correspond to probability models"

Martin C. Frith

2019-03-18

#### 1 Efficient algorithms for local alignment

As described previously [ZPM97, CWC04], Algorithm II can be made more efficient by using these variables:

$$\hat{Y}_{ij} = Y_{i+1\ j}$$
$$\hat{Z}_{ij} = Z_{i\ j+1}$$

The algorithm can then be performed as follows:

**Algorithm S.I**

$$\begin{aligned} W_{00} &= 0 && && \\ W_{i\,0} &= 0 && \hat{Z}_{i\,0} = -\infty && \forall_{i=1}^{m} \\ W_{0\,j} &= 0 && \hat{Y}_{0\,j} = -\infty && \forall_{j=1}^{n} \end{aligned}$$

$$\left.\begin{aligned} W_{ij} &= \max(W_{i-1\ j-1} + S_{R_i Q_j},\ \hat{Y}_{i-1\ j},\ \hat{Z}_{i\ j-1},\ 0) \\ \hat{Y}_{ij} &= \max(W_{ij} + a_D,\ \hat{Y}_{i-1\ j} + b_D) \\ \hat{Z}_{ij} &= \max(W_{ij} + a_I,\ \hat{Z}_{i\ j-1} + b_I) \end{aligned}\right\} \forall_{i=1}^{m}\ \forall_{j=1}^{n}$$

$$\text{Optimal alignment score} = \max_{i=0}^{m} \max_{j=0}^{n} W_{ij}$$

The main advantages are that this algorithm uses only three old values of $W$, $\hat{Y}$, and $\hat{Z}$, whereas Algorithm II uses five old values, and if $a_D = a_I$ (as is often the case) one addition is saved. A further nuance is described in [CWC04].

##### 1.1 Alternatives

An equally-efficient alternative is to use these variables:

$$\check{Y}_{ij} = Y_{i+1\ j} - b_D$$
$$\check{Z}_{ij} = Z_{i\ j+1} - b_I$$

Another possibility is to use $\hat{X}_{ij} = X_{i+1\ j+1}$. These changes of variables can also be used to accelerate the Forward algorithms described in this study.

In practice, a fast implementation should consider processor hardware details in order to optimize the speed and parallelism of calculation [Far07, Rog11, SK18]. Some aspects of this are quite general, e.g. maximizing parallelism by minimizing the number of bits needed to hold each value, and minimizing critical path length [SK18].

### 2 More on the linear-gap model

This section provides more information on the simple gapped PHMM in Figure 4 of the main text, in particular, an alternative explanation of balanced length probability.

#### 2.1 Forward algorithm

The total probability of all possible alignments can be calculated like this:

**Algorithm S.II**

$$\begin{aligned} W'_{00} &= 1 \\ W'_{i\,0} &= W'_{i-1\ 0} \cdot a'_D + 1 && \forall_{i=1}^{m} \\ W'_{0\,j} &= W'_{0\ j-1} \cdot a'_I + 1 && \forall_{j=1}^{n} \\ W'_{ij} &= W'_{i-1\ j-1} \cdot S'_{R_i Q_j} + W'_{i-1\ j} \cdot a'_D + W'_{i\ j-1} \cdot a'_I + 1 && \forall_{i=1}^{m} \forall_{j=1}^{n} \end{aligned}$$

$$\sum_{\text{paths}} \text{prob}(\text{path}, R, Q)/\mu_G = \sum_{i=0}^{m} \sum_{j=0}^{n} W'_{ij}$$

#### 2.2 Conserved probability ratio

Let us consider a different explanation of balanced length probability. The idea is to have transition probabilities (arrow probabilities) that favor neither longer nor shorter alignments. For the gapless model (Figure 2), this means: $\gamma = \omega_D \omega_I$. Intuitively, for the linear-gap model, it seems that $\gamma$ should be less than $\omega_D \omega_I$, because an alignment can be extended in alternative ways (with gaps).

Precisely, this idea means conservation of transition probability ratio relative to a null alignment. Algorithm S.II outputs a probability ratio, where the denominator ($\mu_G$) is the probability of a null alignment. The algorithm calculates this from probability ratios of sequence prefixes. In the algorithm, $W'_{ij}$ supplies probability ratio to $W'_{i+1\ j}$, $W'_{i\ j+1}$, and $W'_{i+1\ j+1}$. Specifically, it supplies:

- probability ratio $W'_{ij} \cdot a'_D$ to $W'_{i+1\ j}$,
- probability ratio $W'_{ij} \cdot a'_I$ to $W'_{i\ j+1}$,
- transition probability ratio $W'_{ij} \cdot c'$ to $W'_{i+1\ j+1}$.

Conservation means that the total probability ratio emanating from $W'_{ij}$ neither increases nor decreases:

$$W'_{ij} \cdot c' + W'_{ij} \cdot a'_D + W'_{ij} \cdot a'_I = W'_{ij},$$

which means:

$$c' + a'_D + a'_I = 1.$$

This is equivalent to Equation 27 in [YH01].

### 3 More on affine-gap model A

#### 3.1 Ambiguity and alignment scores

As explained in the main text, model A (Figure 5A) is ambiguous, meaning that different paths through the PHMM yield indistinguishable alignments. In the main text we assumed that "alignment" means "path", and described probabilities and scores of alignments defined in this way. However, it is also possible to define an alignment to be a set of indistinguishable paths.

Let us explore the case where consecutive deletions of lengths $d_1$ and $d_2$, and a single deletion of length $d_1 + d_2$, are regarded as the same alignment (and likewise for insertions). The probability of a length-$d$ deletion starting just after $R_c$ is now:

$$\text{prob}(\text{del}(c, d)) = \alpha_D(\beta_D + \alpha_D(1 - \beta_D))^{d-1}(1 - \beta_D) \prod_{k=1}^{d} \phi_{R_{c+k}}$$

Likewise, the probability of a length-$d$ insertion starting just after $Q_c$ is:

$$\text{prob}(\text{ins}(c, d)) = \alpha_I(\beta_I + \alpha_I(1 - \beta_I))^{d-1}(1 - \beta_I) \prod_{k=1}^{d} \psi_{Q_{c+k}}$$

Therefore, the gap score parameters in this case are:

$$\begin{aligned}
\bar{a}_D &= t \ln\left(\frac{\alpha_D(1 - \beta_D)}{\omega_D}\right) &&= a_D \\
\bar{b}_D &= t \ln\left(\frac{\beta_D + \alpha_D(1 - \beta_D)}{\omega_D}\right) &&\neq b_D \\
\bar{a}_I &= t \ln\left(\frac{\alpha_I(1 - \beta_I)}{\omega_I}\right) &&= a_I \\
\bar{b}_I &= t \ln\left(\frac{\beta_I + \alpha_I(1 - \beta_I)}{\omega_I}\right) &&\neq b_I
\end{aligned}$$

The practical consequence is that scores for gaps of length $> 1$ are less negative, compared to the case where alignment = path. Furthermore, the optimal path might not be a member of the optimal alignment.

If we wish to reverse-engineer a PHMM from score parameters, then we need to decide whether the score parameters apply to paths or to alignments (sets of paths). In the latter case, we could at first calculate the path parameters:

$$\begin{aligned}
b_D &= t \ln(\exp(\bar{b}_D/t) - \exp(\bar{a}_D/t)) \\
b_I &= t \ln(\exp(\bar{b}_I/t) - \exp(\bar{a}_I/t))
\end{aligned}$$

Once we know the path parameters, the situation is the same as in the main text. Some examples suggest that the difference between gap scores for paths and for alignments is typically small (Table S1).

**Table S1:** Examples of gap scores for paths and for sets of indistinguishable paths.

| Matrix | $t$ | $a_D = a_I$ | $b_D = b_I$ | $\bar{b}_D = \bar{b}_I$ |
|---|---|---|---|---|
| HoxD70 | 96.1735 | 430 | 30 | 28.509 |
| | | | 31.514 | 30 |
| Simple | 0.910239 | 8 | 1 | 0.99958 |
| | | | 1.00042 | 1 |
| BLOSUM50 | 4.2199 | 12 | 2 | 1.6228 |
| | | | 2.41427 | 2 |

##### 3.1.1 Linear gap costs and affine parameterization

These gap score parameters have some properties worth noting. On one hand, there is a convenient guarantee: $\bar{a}_D \leq \bar{b}_D$ and $\bar{a}_I \leq \bar{b}_I$. On the other hand, the linear gap cost case $\bar{a}_{D/I} = \bar{b}_{D/I}$ corresponds to $b_{D/I} = -\infty$ and $b'_{D/I} = 0$.

Therefore, it is desirable for the Forward algorithm to allow $b'_{D/I} = 0$. The Forward algorithm presented here (Algorithm III in the main text) does allow this, but only because it uses the right parameterization of affine gaps. The two common parameterizations of affine score for a gap of length $k$ are: $a + b \times (k-1)$ and $a + b \times k$. Algorithm III is based on the former, whereas the latter does not support $b = -\infty$.

##### 3.1.2 Further ambiguity

There is another potential ambiguity in model A: an insertion followed by a deletion might be regarded as indistinguishable from a deletion followed by an insertion. However, if we define these as being the same alignment, then alignment scores do not have the standard affine-gap form (consecutive inserts and deletes score less negatively than non-consecutive ones).

#### 3.2 Conserved probability ratio

Similarly to the linear-gap model, balanced length probability can be understood as conservation of transition probability ratio relative to a null alignment. The Forward algorithm (Algorithm III in the main text) outputs a probability ratio, where the denominator ($\mu_G$) is the probability of a null alignment. The algorithm calculates this from probability ratios of sequence prefixes. Conservation is easier to see if we rescale $Y'$ and $Z'$:

$$Y''_{ij} = Y'_{ij}/(1 - b'_D) \qquad Z''_{ij} = Z'_{ij}/(1 - b'_I)$$

The recurrences in Algorithm III can be written in terms of $Y''$ and $Z''$:

$$\begin{aligned}
W'_{ij} &= X'_{ij} + (1 - b'_D)Y''_{ij} + (1 - b'_I)Z''_{ij} + 1 && \forall_{i=0}^{m} \forall_{j=0}^{n} \\
X'_{ij} &= W'_{i-1\ j-1} \cdot S'_{R_i Q_j} && \forall_{i=1}^{m} \forall_{j=1}^{n} \\
Y''_{ij} &= W'_{i-1\ j} \cdot a'_D/(1 - b'_D) + Y''_{i-1\ j} \cdot b'_D && \forall_{i=1}^{m} \forall_{j=0}^{n} \\
Z''_{ij} &= W'_{i\ j-1} \cdot a'_I/(1 - b'_I) + Z''_{i\ j-1} \cdot b'_I && \forall_{i=0}^{m} \forall_{j=1}^{n}
\end{aligned}$$

The point of this rescaling is that the total probability ratio emanating from $Y''_{ij}$ and $Z''_{ij}$ neither increases nor decreases. The probability ratio emanating from $Y''_{ij}$ is $(1 - b'_D)Y''_{ij}$ (to $W'_{ij}$) and $b'_D Y''_{ij}$ (to $Y''_{i+1\ j}$), which sums to $Y''_{ij}$. Likewise for $Z''_{ij}$. The probability ratio (of transition probabilities) emanating from $W'_{ij}$ is:

$$\left(c' + \frac{a'_D}{1 - b'_D} + \frac{a'_I}{1 - b'_I}\right) W'_{ij}$$

Conservation of probability ratio means that this should equal $W'_{ij}$. Equivalently:

$$\frac{\gamma}{\omega_D \omega_I} + \frac{\alpha_D(1 - \beta_D)}{\omega_D - \beta_D} + \frac{\alpha_I(1 - \beta_I)}{\omega_I - \beta_I} = 1$$

### 4 Model B

This section deals with the PHMM in Figure 5B of the main text. The math turns out to become simpler if we use the M→M transition probability, $\Gamma = \gamma(1 - \alpha_D - \alpha_I)$, instead of $\gamma$.

#### 4.1 Constraint on $\delta$

If we desire standard affine-gap alignment, $\delta$ should be set such that adjacent deletions and insertions have the same probability as non-adjacent deletions and insertions. That is, these two configurations should have the same probability:

```
OO-OO   OOO-O
O-OOO   O-OOO
```

This means that:

$$\text{prob}(D \to I) \cdot \text{prob}(M \to M) = \text{prob}(D \to M) \cdot \text{prob}(M \to I),$$

which implies that:

$$\delta = \frac{(1 - \beta_D)\alpha_I}{\Gamma + \alpha_I}$$

#### 4.2 Scores in terms of model probabilities

Consider an alignment where the first $c$ letters of $R$ and $Q$ are aligned, the next $d$ letters of $R$ are inserted, the next $e$ letters of $R$ and $Q$ are aligned, the next $f$ letters of $Q$ are inserted, and the final $g$ letters of $R$ and $Q$ are aligned. The probability of this alignment under the model is:

$$\begin{aligned}
\text{prob}(\text{path}, R, Q) = {} & (1-\omega_D)(1-\omega_I) \\
& \left(\prod_{k=1}^{c} \pi_{R_k Q_k}\right) \gamma^c (1-\alpha_D-\alpha_I)^{c-1} \\
& \left(\prod_{k=1}^{d} \phi_{R_{c+k}}\right) \alpha_D \beta_D^{d-1} (1-\beta_D-\delta) \\
& \left(\prod_{k=1}^{e} \pi_{R_{c+d+k} Q_{c+k}}\right) \gamma^{e-1} (1-\alpha_D-\alpha_I)^{e-1} \\
& \left(\prod_{k=1}^{f} \psi_{Q_{c+e+k}}\right) \alpha_I \beta_I^{f-1} (1-\beta_I) \\
& \left(\prod_{k=1}^{g} \pi_{R_{c+d+e+k} Q_{c+e+f+k}}\right) \gamma^{g-1} (1-\alpha_D-\alpha_I)^{g} \\
& (1-\gamma)(1-\omega_D)(1-\omega_I)
\end{aligned}$$

We can rearrange this to:

$$\begin{aligned}
\text{prob}(\text{path}, R, Q) = {} & (1-\omega_D)^2 (1-\omega_I)^2 (1-\gamma) \\
& \prod_{k=1}^{c} \Gamma \pi_{R_k Q_k} \\
& \frac{\alpha_D(1-\beta_D-\delta)}{\Gamma} \beta_D^{d-1} \prod_{k=1}^{d} \phi_{R_{c+k}} \\
& \prod_{k=1}^{e} \Gamma \pi_{R_{c+d+k} Q_{c+k}} \\
& \frac{\alpha_I(1-\beta_I)}{\Gamma} \beta_I^{f-1} \prod_{k=1}^{f} \psi_{Q_{c+e+k}} \\
& \prod_{k=1}^{g} \Gamma \pi_{R_{c+d+e+k} Q_{c+e+f+k}}
\end{aligned}$$

Dividing by $\mu$ (defined in the main text) gives:

$$\begin{aligned}
\frac{\text{prob}(\text{path}, R, Q)}{\mu} = {} & \prod_{k=1}^{c} \frac{\Gamma}{\omega_D \omega_I} \frac{\pi_{R_k Q_k}}{\phi_{R_k} \psi_{Q_k}} \\
& \frac{\alpha_D(1-\beta_D-\delta)}{\Gamma \omega_D} \left(\frac{\beta_D}{\omega_D}\right)^{d-1} \\
& \prod_{k=1}^{e} \frac{\Gamma}{\omega_D \omega_I} \frac{\pi_{R_{c+d+k} Q_{c+k}}}{\phi_{R_{c+d+k}} \psi_{Q_{c+k}}} \\
& \frac{\alpha_I(1-\beta_I)}{\Gamma \omega_I} \left(\frac{\beta_I}{\omega_I}\right)^{f-1} \\
& \prod_{k=1}^{g} \frac{\Gamma}{\omega_D \omega_I} \frac{\pi_{R_{c+d+e+k} Q_{c+e+f+k}}}{\phi_{R_{c+d+e+k}} \psi_{Q_{c+e+f+k}}}
\end{aligned}$$

Now we can express the score parameters in terms of the model probabilities:

$$
\begin{aligned}
S_{xy} &= t \ln\left(\frac{\pi_{xy}}{\phi_x \psi_y} \cdot \frac{\Gamma}{\omega_D \omega_I}\right) \\
a_D &= t \ln\left(\frac{\alpha_D(1-\beta_D-\delta)}{\Gamma \omega_D}\right) \\
b_D &= t \ln\left(\frac{\beta_D}{\omega_D}\right) \\
a_I &= t \ln\left(\frac{\alpha_I(1-\beta_I)}{\Gamma \omega_I}\right) \\
b_I &= t \ln\left(\frac{\beta_I}{\omega_I}\right)
\end{aligned}
$$

Finally, substituting in the constrained value of $\delta$ gives:

$$
a_D = t \ln\left(\frac{\alpha_D(1-\beta_D)}{(\Gamma+\alpha_I)\omega_D}\right)
$$

#### 4.3 Symmetric insertions and deletions

Suppose we wish to make the model as symmetric as possible, and so we specify that $\omega_D = \omega_I$, $\beta_D = \beta_I$, and $a_D = a_I$. In that case, $\alpha_D$ and $\alpha_I$ are related like this:

$$
\alpha_D = \alpha_I + \alpha_I^2/\Gamma
$$

#### 4.4 Limits to degrees of freedom

Model B has several degrees of freedom, meaning that we can vary the model probabilities with no effect on alignment scores. In particular, we can freely vary $\omega_D$ and $\omega_I$, provided we co-vary $\beta_D$, $\beta_I$, $\alpha_D$, $\alpha_I$, and $\gamma$ so as to keep these values fixed:

$$
\begin{aligned}
c' &= \Gamma/(\omega_D \omega_I) \\
a'_D &= \exp(a_D/t) = \frac{\alpha_D(1-\beta_D)}{(\Gamma+\alpha_I)\omega_D} \\
b'_D &= \exp(b_D/t) = \beta_D/\omega_D \\
a'_I &= \exp(a_I/t) = \frac{\alpha_I(1-\beta_I)}{\Gamma \omega_I} \\
b'_I &= \exp(b_I/t) = \beta_I/\omega_I
\end{aligned}
$$

We cannot, however, have $\gamma \geq 1$. Thus:

$$
\gamma = \Gamma/(1-\alpha_D-\alpha_I) < 1
$$

Equivalently:

$$
\Gamma + \alpha_D + \alpha_I < 1
$$

We can express $\Gamma + \alpha_D + \alpha_I$ in terms of $\omega_D$, $\omega_I$, and the fixed values, as follows:

$$\alpha_I = c'\omega_D\omega_I \cdot \frac{a'_I\omega_I}{1 - b'_I\omega_I}$$

$$\alpha_D = c'\omega_D\omega_I \cdot \frac{a'_D\omega_D(1 - b'_I\omega_I + a'_I\omega_I)}{(1 - b'_D\omega_D)(1 - b'_I\omega_I)}$$

$$\Gamma + \alpha_D + \alpha_I = c'\omega_D\omega_I \cdot \frac{(1 - b'_D\omega_D + a'_D\omega_D)(1 - b'_I\omega_I + a'_I\omega_I)}{(1 - b'_D\omega_D)(1 - b'_I\omega_I)}$$

Thus, $\omega_D$ and $\omega_I$ are restricted to values that make the right-hand side of the above equation $< 1$.

#### 4.5 Balanced length probability

Balanced length probability occurs when:

$$c' \cdot \frac{(1 - b'_D + a'_D)(1 - b'_I + a'_I)}{(1 - b'_D)(1 - b'_I)} = 1$$

In this case, we can freely vary $\omega_D$, $\omega_I$, and $\gamma$ arbitrarily close to 1. This is equivalent to Equation 28 in [YH01].

#### 4.6 Forward algorithm

The Forward algorithm for model B can be expressed like this:

**Algorithm S.III**

$$X'_{i\,0} = Y'_{i\,0} = Z'_{i\,0} = 0 \qquad \forall_{i=0}^{m}$$
$$X'_{0\,j} = Y'_{0\,j} = Z'_{0\,j} = 0 \qquad \forall_{j=0}^{n}$$

$$\begin{aligned}
W'_{ij} &= X'_{ij} + 1 && \forall_{i=0}^{m}\forall_{j=0}^{n} \\
X'_{ij} &= (Y'_{i-1\;j-1} + Z'_{i-1\;j-1} + W'_{i-1\;j-1}) \cdot S'_{R_iQ_j} && \forall_{i=1}^{m}\forall_{j=1}^{n} \\
Y'_{ij} &= X'_{i-1\;j} \cdot a'_D + Y'_{i-1\;j} \cdot b'_D && \forall_{i=1}^{m}\forall_{j=1}^{n} \\
Z'_{ij} &= X'_{i\;j-1} \cdot a'_I + Y'_{i\;j-1} \cdot a'_I + Z'_{i\;j-1} \cdot b'_I && \forall_{i=1}^{m}\forall_{j=1}^{n}
\end{aligned}$$

$$\frac{\mathrm{prob}(R, Q)}{\mu} = \sum_{i=0}^{m}\sum_{j=0}^{n} W'_{ij}$$

#### 4.7 Conserved probability ratio

Let us consider transition probabilities that conserve probability ratio relative to a null alignment. Similarly to model A, conservation is easier to see if we rescale $Y'$ and $Z'$:

$$Z''_{ij} = Z'_{ij} \cdot c' \cdot \frac{1}{1 - b'_I}$$

$$Y''_{ij} = Y'_{ij} \cdot c' \cdot \frac{1 - b'_I + a'_I}{(1 - b'_D)(1 - b'_I)}$$

If the Forward algorithm is written in terms of $Y''$ and $Z''$, it can be verified that the total probability ratio (of transition probabilities) emanating from $Y''_{ij}$ and $Z''_{ij}$ neither increases nor decreases. The probability ratio emanating from $X'_{ij}$ is:

$$c' \cdot \frac{(1 - b'_D + a'_D)(1 - b'_I + a'_I)}{(1 - b'_D)(1 - b'_I)} \cdot X'_{ij}$$

Conservation of probability ratio means that this should equal $X'_{ij}$. Equivalently:

$$\left(\Gamma + \alpha_I + \frac{1 - \beta_D}{\omega_D - \beta_D} \cdot \alpha_D\right)\left(1 + \frac{1 - \omega_I}{\omega_I - \beta_I} \cdot \frac{\alpha_I}{\Gamma + \alpha_I}\right) = \omega_D \omega_I$$

Models that conserve probability ratio are precisely those with balanced length probability.

#### 4.8 Maximum likelihood parameters

A standard way of selecting PHMM parameters is by seeking a maximum-likelihood fit to some training data [DEKM98]. This procedure (i) calculates the expected count of each transition and emission, and (ii) selects model parameters that maximize the likelihood of those counts. Step (ii) is affected if we impose constraints on the parameters (such as the constraint on $\delta$), so let us examine step (ii) here.

Suppose we are given these expected counts:

$c_m$**:** matches plus mismatches

$c_D$**:** deleted letters

$c_I$**:** inserted letters

$c_d$**:** runs of deleted letters

$c_i$**:** runs of inserted letters

$c_p$**:** adjacent pairs of insertions and deletions

The probability of observing these counts under the model is proportional to:

$$\begin{aligned}
&\beta_I^{c_I - c_i} \cdot (1 - \beta_I)^{c_i} \\
&\beta_D^{c_D - c_d} \cdot (1 - \beta_D - \delta)^{c_d - c_p} \cdot \delta^{c_p} \\
&\alpha_D^{c_d} \cdot \alpha_I^{c_i - c_p} \cdot (1 - \alpha_D - \alpha_I)^{c_m - c_d - c_i + c_p} \\
&\gamma^{c_m - c_d - c_i + c_p} \cdot (1 - \gamma)
\end{aligned}$$

We wish to select parameters (Greek letters) that maximize this probability, or equivalently, maximize its logarithm:

$$\begin{aligned}
&(c_I - c_i) \ln \beta_I + c_i \ln(1 - \beta_I) \\
&+ (c_D - c_d) \ln \beta_D + (c_d - c_p) \ln(1 - \beta_D - \delta) + c_p \ln \delta \\
&+ c_d \ln \alpha_D + (c_i - c_p) \ln \alpha_I + (c_m - c_d - c_i + c_p) \ln(1 - \alpha_D - \alpha_I) \\
&+ (c_m - c_d - c_i + c_p) \ln \gamma + \ln(1 - \gamma)
\end{aligned}$$

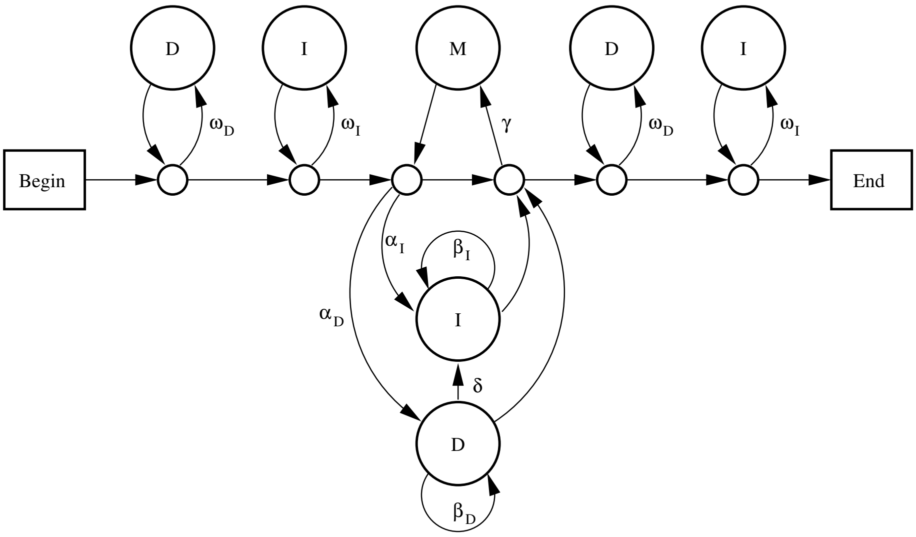


**Figure S1:** Model C for gapped local alignment. The state labeled M emits aligned letters $x : y$ with probability $\pi_{xy}$. States labeled D emit reference letters $x$ with probability $\phi_x$. States labeled I emit query letters $y$ with probability $\psi_y$.

Substituting in the constrained value of $\delta$ gives:

$$
\begin{aligned}
&(c_I - c_i)\ln\beta_I + c_i\ln(1-\beta_I)\\
&+ (c_D - c_d)\ln\beta_D + c_d\ln(1-\beta_D)\\
&- c_d\ln(\gamma(1-\alpha_D-\alpha_I)+\alpha_I)\\
&+ c_d\ln\alpha_D + c_i\ln\alpha_I + (c_m - c_i)\ln(1-\alpha_D-\alpha_I)\\
&+ (c_m - c_i)\ln\gamma + \ln(1-\gamma)
\end{aligned}
$$

Without the constraint, the maximum-likelihood parameters are straightforward:

$$
\begin{aligned}
\alpha_D &= c_d/c_m\\
\alpha_I &= (c_i - c_p)/c_m\\
\beta_D &= (c_D - c_d)/c_D\\
\beta_I &= (c_I - c_i)/c_I\\
\gamma &= (c_m - c_d - c_i + c_p)/(c_m - c_d - c_i + c_p + 1)\\
\delta &= c_p/c_D
\end{aligned}
$$

With the constraint, the maximum-likelihood $\beta_D$ and $\beta_I$ are unchanged, but unfortunately $\alpha_D$, $\alpha_I$, and $\gamma$ do not appear to be simple.

### 5 Yet another model of gapped local alignment

Model C (Figure S1) is yet another model of gapped local alignment: this one is considered here because it produces recurrence relations of the form in [YH01].

#### 5.1 Constraint on $\delta$

If we desire standard affine-gap alignment, $\delta$ should be set such that adjacent deletions and insertions have the same probability as non-adjacent deletions and insertions. By similar reasoning to that used above for model B:

$$\delta = \frac{(1-\beta_D)\alpha_I}{1-\alpha_D}$$

#### 5.2 Scores in terms of model probabilities

Similarly to the other PHMMs, the alignment probabilities can be connected to scores. First, define $\mu_C = \text{prob}(\text{null path}, R, Q)$, where "null path" means a path that never uses the $\alpha_D$, $\alpha_I$, or $\gamma$ arrows:

$$\mu_C = \left(\prod_{k=1}^{m} \omega_D \phi_{R_k}\right)\left(\prod_{k=1}^{n} \omega_I \psi_{Q_k}\right)(1-\omega_D)^2(1-\omega_I)^2(1-\gamma)(1-\alpha_D-\alpha_I)$$

Then, for any path, $t\ln(\text{prob}(\text{path}, R, Q)/\mu_C)$ is a sum of these affine-gap scores:

$$S_{xy} = t\ln\left(\frac{\pi_{xy}}{\phi_x\psi_y}\cdot\frac{\Gamma}{\omega_D\omega_I}\right)$$

$$a_D = t\ln\left(\frac{\alpha_D(1-\beta_D)}{\omega_D(1-\alpha_D)}\right)$$

$$b_D = t\ln\left(\frac{\beta_D}{\omega_D}\right)$$

$$a_I = t\ln\left(\frac{\alpha_I(1-\beta_I)}{\omega_I(1-\alpha_D-\alpha_I)}\right)$$

$$b_I = t\ln\left(\frac{\beta_I}{\omega_I}\right)$$

#### 5.3 Symmetric insertions and deletions

Suppose we wish to make the model as symmetric as possible, and so we specify that $\omega_D = \omega_I$, $\beta_D = \beta_I$, and $a_D = a_I$. In that case, $\alpha_D$ and $\alpha_I$ are related like this:

$$\alpha_I = \alpha_D(1-\alpha_D)$$

#### 5.4 Limits to degrees of freedom

Model C has several degrees of freedom, meaning that we can vary the model probabilities with no effect on alignment scores. In particular, we can freely vary $\omega_D$ and $\omega_I$, provided we co-vary $\beta_D$, $\beta_I$, $\alpha_D$, $\alpha_I$, and $\gamma$ so as to keep these values fixed:

$$\begin{aligned} c' &= \Gamma/(\omega_D \omega_I) \\ a'_D &= \exp(a_D/t) = \frac{\alpha_D(1-\beta_D)}{\omega_D(1-\alpha_D)} \\ b'_D &= \exp(b_D/t) = \beta_D/\omega_D \\ a'_I &= \exp(a_I/t) \ = \frac{\alpha_I(1-\beta_I)}{\omega_I(1-\alpha_D-\alpha_I)} \\ b'_I &= \exp(b_I/t) \ = \beta_I/\omega_I \end{aligned}$$

We cannot, however, have $\gamma \geq 1$. We can express $\gamma$ in terms of $\omega_D$, $\omega_I$, and the fixed values, as follows:

$$\begin{aligned} \alpha_D &= \frac{a'_D \omega_D}{1 - b'_D \omega_D + a'_D \omega_D} \\ \alpha_I &= \frac{a'_I \omega_I (1 - b'_D \omega_D)}{(1 - b'_D \omega_D + a'_D \omega_D)(1 - b'_I \omega_I + a'_I \omega_I)} \\ 1 - \alpha_D - \alpha_I &= \frac{(1 - b'_D \omega_D)(1 - b'_I \omega_I)}{(1 - b'_D \omega_D + a'_D \omega_D)(1 - b'_I \omega_I + a'_I \omega_I)} \\ \gamma &= c' \omega_D \omega_I \cdot \frac{(1 - b'_D \omega_D + a'_D \omega_D)(1 - b'_I \omega_I + a'_I \omega_I)}{(1 - b'_D \omega_D)(1 - b'_I \omega_I)} \end{aligned}$$

Thus, $\omega_D$ and $\omega_I$ are restricted to values that make the right-hand side of the above equation $< 1$. This turns out to be the same restriction as for model B!

#### 5.5 Forward algorithm

The Forward algorithm for model C can be expressed like this:

**Algorithm S.IV**

$$\begin{aligned} X'_{i\,0} &= 1 \qquad Z'_{i\,0} = 0 \qquad \forall_{i=0}^{m} \\ X'_{0\,j} &= 1 \qquad Y'_{0\,j} = 0 \qquad \forall_{j=0}^{n} \end{aligned}$$

$$\begin{aligned} W'_{ij} &= X'_{ij} + Y'_{ij} + Z'_{ij} & \forall_{i=0}^{m} \forall_{j=0}^{n} \\ X'_{ij} &= W'_{i-1\ j-1} \cdot S'_{R_i Q_j} + 1 & \forall_{i=1}^{m} \forall_{j=1}^{n} \\ Y'_{ij} &= X'_{i-1\ j} \cdot a'_D + Y'_{i-1\ j} \cdot b'_D & \forall_{i=1}^{m} \forall_{j=0}^{n} \\ Z'_{ij} &= X'_{i\ j-1} \cdot a'_I + Y'_{i\ j-1} \cdot a'_I + Z'_{i\ j-1} \cdot b'_I & \forall_{i=0}^{m} \forall_{j=1}^{n} \end{aligned}$$

$$\frac{\text{prob}(R,Q)}{\mu_C} = \sum_{i=0}^{m} \sum_{j=0}^{n} W'_{ij}$$

#### 5.6 Conserved probability ratio

The transition probabilities of model C conserve probability ratio in the same situation as model B (because the Forward algorithms are similar):

$$c' \cdot \frac{(1 - b'_D + a'_D)(1 - b'_I + a'_I)}{(1 - b'_D)(1 - b'_I)} = 1$$

For model C, this is equivalent to:

$$\frac{\gamma}{\omega_D \omega_I}\left(1+\frac{1-\omega_D}{\omega_D-\beta_D}\cdot\alpha_D\right)\left(1+\frac{1-\omega_I}{\omega_I-\beta_I}\cdot\frac{\alpha_I}{1-\alpha_D}\right)=1$$

Again, parameter sets that conserve probability ratio are those with balanced length probability (where we can freely vary $\omega_D$, $\omega_I$, and $\gamma$ arbitrarily close to 1).

##### 5.7 Maximum likelihood parameters

It is useful to select PHMM parameters that have maximum-likelihood fit to some given sequence data. As mentioned above for model B, this involves finding model parameters that maximize the likelihood of some expected counts ($c_m, c_D, c_I, c_d, c_i, c_p$, defined above). The probability of observing these counts under the model is proportional to:

$$\begin{aligned}
&\beta_I^{c_I-c_i}\cdot(1-\beta_I)^{c_i}\\
&\beta_D^{c_D-c_d}\cdot(1-\beta_D-\delta)^{c_d-c_p}\cdot\delta^{c_p}\\
&\alpha_D^{c_d}\cdot\alpha_I^{c_i-c_p}\cdot(1-\alpha_D-\alpha_I)^{c_m-c_d-c_i+c_p+1}\\
&\gamma^{c_m}\cdot(1-\gamma)
\end{aligned}$$

We wish to select parameters (Greek letters) that maximize this probability, or equivalently, maximize its logarithm:

$$\begin{aligned}
&(c_I-c_i)\ln\beta_I+c_i\ln(1-\beta_I)\\
&+(c_D-c_d)\ln\beta_D+(c_d-c_p)\ln(1-\beta_D-\delta)+c_p\ln\delta\\
&+c_d\ln\alpha_D+(c_i-c_p)\ln\alpha_I+(c_m-c_d-c_i+c_p+1)\ln(1-\alpha_D-\alpha_I)\\
&+c_m\ln\gamma+\ln(1-\gamma)
\end{aligned}$$

Substituting in the constrained value of $\delta$ gives:

$$\begin{aligned}
&(c_I-c_i)\ln\beta_I+c_i\ln(1-\beta_I)\\
&+(c_D-c_d)\ln\beta_D+c_d\ln(1-\beta_D)\\
&-c_d\ln(1-\alpha_D)+c_d\ln\alpha_D+c_i\ln\alpha_I+(c_m-c_i+1)\ln(1-\alpha_D-\alpha_I)\\
&+c_m\ln\gamma+\ln(1-\gamma)
\end{aligned}$$

Then, the maximum likelihood parameters are:

$$\begin{aligned}
\alpha_D&=c_d/(c_m+1)\\
\alpha_I&=c_i(c_m-c_d+1)/(c_m+1)^2\\
\beta_D&=(c_D-c_d)/c_D\\
\beta_I&=(c_I-c_i)/c_I\\
\gamma&=c_m/(c_m+1)
\end{aligned}$$

[DEKM98] Richard Durbin, Sean Eddy, Anders Krogh, and Graeme Mitchison. *Biological sequence analysis: probabilistic models of proteins and nucleic acids.* Cambridge University Press, 1998.

[Far07] M. Farrar. Striped Smith-Waterman speeds database searches six times over other SIMD implementations. *Bioinformatics*, 23(2):156–161, Jan 2007.

[Rog11] T. Rognes. Faster Smith-Waterman database searches with inter-sequence SIMD parallelisation. *BMC Bioinformatics*, 12:221, 2011.

[SK18] H. Suzuki and M. Kasahara. Introducing difference recurrence relations for faster semi-global alignment of long sequences. *BMC Bioinformatics*, 19(Suppl 1):45, Feb 2018.

[YH01] Y. K. Yu and T. Hwa. Statistical significance of probabilistic sequence alignment and related local hidden Markov models. *J. Comput. Biol.*, 8(3):249–282, 2001.

[ZPM97] Z. Zhang, W. R. Pearson, and W. Miller. Aligning a DNA sequence with a protein sequence. *J. Comput. Biol.*, 4(3):339–349, 1997.
